# Distance v.s. Resolution: Neuromapping of Effective Resolution onto Physical Distance

**DOI:** 10.1101/2023.08.03.551725

**Authors:** Suayb S. Arslan, Michal Fux, Pawan Sinha

## Abstract

The main focus of this work is on determining the effective resolution of a face image on the retina when the face is at a particular distance from the eye. Despite its straightforward articulation, arriving at a satisfactory solution might be unexpectedly challenging. The relationship between viewing distance and effective resolution cannot be easily obtained through Snellen acuity, contrast sensitivity, photoreceptor packing density or ganglion cell convergence rate in the retina. We used theoretical modelling to establish preliminary guidelines and then tested them empirically. We showed participants a 2 *×* 2 array of images in different resolutions at various viewing distances. At each distance, participants were expected to perform an “odd one out” task, identifying the image that was blurrier than the other three. As the study progressed, viewing distance was gradually reduced. The data collected enabled us to determine the upper and lower limits of the available effective resolution for human vision under normal conditions, as a function of viewing distance. Notably, human performance in blur detection is superior to what a theoretical model based on projected image size, cone density, and foveal extent predicts, especially at close ranges. Therefore, we propose that future theoretical models must account for non-uniform in–fovea density and the less pronounced decline in acuity outside the fovea to establish a reliable relationship between viewing distance and perceived image characteristics. The <distance:effective-resolution> mapping allows for a direct comparison of human face recognition performance across different levels of blur and viewing distance. It also enables us to systematically compare human performance to that of machine vision systems using resolution as a common factor.

## 1. Introduction

A practical challenge arises in characterizing information due to viewing conditions at varying distances, particularly when considering the diverse front ends of distinct vision systems. It is fundamental to study this unaddressed challenge in a manner that would allow for translation across different vision systems, each with their unique optical properties. By doing so, we not only tackle the fundamental limitations of visual perception in space but also create an opportunity to test theoretical models of information sampling. An effective solution to this challenge has the potential to open up additional avenues of research, including the impact of scale on information acquisition and subsequent processing steps. This holistic approach promises to deepen our understanding of how visual systems adapt to different viewing conditions and scales as well as help compare humans and machines under the same set of optical frontend assumptions. In addition, having a proven functional relationship between the viewing distance and available image resolution for the human visual system would enable a principled comparison of the human recognition performance with respect to machine performance. This will not only help us understand how much are humans able to see with increasing distances/ranges but also would be conducive to guiding bio-inspired neural network designs taking into account the limitations of the human optical/visual system. Furthermore, addressing the challenges posed by impoverished images and their inherent prevalence in long-range viewing conditions would enable us to implicitly assess the performance of individuals with low vision by appropriately adjusting the distance axis. This valuable information holds significant relevance in the development of rehabilitation programs and assistive devices aimed at improving overall health and well-being. In our study, we shall consider face stimuli for further investigation due to their pivotal role in human communication and interpersonal interaction.

In comparison to up-close viewing, the perception of facial characteristics from a distance presents a new set of challenges, including changes in the relative sizes (reduction in the available resolution) and geometry and orientations of the features as well as the addition of irrelevant background information. Because of these variations, it is important for scientists to develop specific models to describe how the visual system interprets facial information at a distance (1–4). Given the changes in the social environment between close-up and distant viewing, the impact of distance and available spatial resolution on the performance of facial recognition task must also be taken into account. The level of the interaction decreases over a distance, as does the capacity to interpret non-verbal cues. Therefore, it is scientifically interesting to consider how the physical proximity in terms of transferred information content affect people’s capacity for distant face recognition. Particularly relevant is the resultant information bottleneck that transpires when faces are viewed at a distance, which can be quantified in terms of the number of pixels (spatial resolution) that can be represented by the available photoreceptors. Unfortunately, there is no simple/linear relationship between the physiological capacity of human vision and the *effective* resolution that is perceived in a given recognition task. The absence of this mapping appears to be a foundational element that is notably missing when juxtaposing human performance against artificial networks commonly employed in computer vision applications (5, 6).

Despite the shortage of data on viewing faces at a distance, there is helpful data from previous research that has investigated the roles of different spatial frequencies in face recognition, which is particularly relevant to viewing faces from a distance. These investigations have demonstrated that low spatial frequencies permit configural processing, whereas high spatial frequencies assist in featural analysis by providing detailed information at closer viewing ranges (7–10). Due to the anticipated impact of local diffusive filtering in distant viewing conditions, it is expected that the degradation of local featural details will be more pronounced compared to larger scale configural information (11, 12). In fact, it has been shown that reduced image resolution can even improve configural processing, especially when all available resources are dedicated to combining high-level image cues (8). The global visual information in a picture may be made simpler with low spatial frequencies present compared to full resolution, which makes it easier for the brain to infer the connections between various characteristics. It is instrumental to comprehend the context of an image when the details are less apparent because the brain can concentrate on the general structure and connections between the dependent long-range elements. This is quite natural to see in human visual system in which low resolution seems to be essential at the early stages of their development to maintain a successful holistic processing (13).

One plausible hypothesis to explain the improved face recognition performance of humans suggests that under long-range viewing conditions, configural processing may play a crucial role in the recognition process, while featural information could be largely obliterated as a result of local diffusions. However, as the viewing distance diminishes, holistic processing comes into play and is enhanced by the presence of featural cues, thereby yielding higher effective resolutions beyond what can be achieved solely through available photoreceptors. This specific issue forms the focal point of our research, as we aim to explicitly measure the actual available resolution of a face at different distances to quantify the information content available for the human visual system to employ these strategies (14). In a broader context, the findings of our study would improve the direct comparison between deep neural networks and human performance in such visual tasks that involve distance-related degraded conditions, representative of real-life viewing scenarios (15). The prevailing approach, so far, has been employing Gaussian “blur” as the primary axis of quality degradation. However, we propose that introducing an intermediate axis of effective resolution is an integral component, as it encapsulates the human processing mechanisms that map the corresponding level of Gaussian blur to the physical distance.

Fitting an appropriate theoretical curve to the data, based on our distance to effective resolution mapping, requires making anatomical assumptions about the structures and functions that underlie the human visual process. We suggested such a parameterized theoretical model the parameters of which are optimized based on the collected data. In the following sections, we elaborate on these parameters and the assumptions that we took into consideration, for example the distribution of photoreceptor cells in the retina and convergence rate from photoreceptor cells to ganglion cells.

The current study contributes to a growing field of investigation of face perception and recognition under degraded conditions. Past studies have focused on the effects of such conditions as resolution (4) and distance and illumination (3) on face recognition accuracy. For example, Wagenaar and Van Der Schrier have found a systematic increase of recognition performance with decreasing distance and increasing illumination. However, it remains unclear how much of the visual information of the face is actually available for the visual system under those differing viewing conditions such as resolution, distance and illumination. A recent account of human face recognition performance, as a function of distance (16, 17), reveals that the salience distribution across a face changes qualitatively with increasing distance. Through decomposing the distinguishable facial features, they have revealed the significance of relationships between internal facial features and external head contours for recognition as a function of distance, which has implications for both human and artificial visual systems (18).

Only a handful of studies have focused on recognition as a function of range (2, 3, 19). Some of these studies focused specifically on examining the impact of resolution on the featural and configural cues within facial images (20–22), elucidating their consequential influence on recognition performance and underscoring the overarching significance of holistic facial feature processing.

There exists a diverse set of applications that present an ample opportunity to investigate a rich array of research questions related to distance-resolution mapping. These applications include but are not limited to courtrooms and eyewitness identification (2, 23). Additionally, surveillance settings often utilize near-infrared face recognition technology, particularly under-illuminated environments, to recognize faces from a distance (24). Since the resolution of the subtended image on the retina is expected to decrease when the subjects are viewed at a distance, low-resolution face recognition has been paid a particular attention in literature. Particularly, the threshold (minimal) resolution is investigated, above which the performance fairly stays steady and while below which the performance deteriorates rapidly (25, 26). In addition to low quality, various non-linear degradations are reported to be challenging for face recognition performance and hence require a special attention (27). A characteristic shared among previous studies is the utilization of a uniform parameter axis to manipulate degradation, such as scaling or blurring, resulting from remote viewing conditions, while leaving the actual mapping between perceived resolution of the viewed face as a function of distance unexplored.

In fact, the mapping between the viewing angle and distance as well as the effective resolution is not necessarily obvious and may highly be dependent on the athmospheric conditions (haze, smoke, etc.), the subject’s acuity, object perception capabilities and strategies, ventral and dorsal processing (fast/slow) (28). In fact, part of the research objectives of this study is to investigate this relationship under controlled experimental set-ups. Another part of this study is aimed at exploring different strategies which may be a result of integration processes within the human brain, specifically focusing on longer viewing time periods that allow for the perceptual differentiation between similar objects. In other words, as viewing distances increase, individuals tend to allocate more time to integrate multiple snapshots of the same observed optotype, gradually enhancing their effective resolution over time. Conversely, for shorter viewing distances, subjects require less processing time to adapt their optical/visual system and achieve the sufficient effective resolution at a given distance. Analyzing this integration process in terms of latency reveals distinct strategies adapted by humans, manifesting itself in the form of *distinct patterns of failed and correct decisions*. Based on our review over the previous research literature, it appears that a comprehensive theoretical framework is missing that explains the effective resolution v.s. distance mapping as a function of the processing time. It seems that the establishment of any framework would rely heavily on the presence of a robust predictive model, which encompasses accurate modeling of physiological and cognitive processes, as well as a meticulously designed experimental setup to ensure proper validation. Collaboration among experts from diverse fields is crucial to address this gap and advance human-computer interaction and multimedia systems in line with human perceptual and cognitive capabilities.

## 2. Theoretical Estimates of Effective Resolution as a Function of Viewing Distance

### 2.1. Optical limits, acuity and effective resolution

The maximum degree of spatial partitioning discrimination in the eye’s object space is referred to as visual acuity (29). The standard definition of ordinary human visual acuity, also called *letter acuity*, is said to be matched when the foveal resolution reaches a 20/20 Snellen Fraction (SF) (or 6/6 internationally). For instance, an optical system that is able to achieve double this acuity level is described as 20/10 or half of this acuity as 20/40. Acuity can be expressed in terms of cycles per second (cpd) where the cycle period is the minimum distance between the two peaks of a periodic signal (typically sinusoidal) with high contrast. Foveal acuity (*F*) is simply a linear function of the SF and can be given as *F* = 30 cpd *×* SF.

In our work, the receptive field was partitioned into two distinct regions: the photoreceptors inside the fovea (IF) and those outside the fovea (OF). To maintain simplicity, our analysis solely focused on the cones (with the minimal assumed impact of rods). Notably, when it comes to long-range viewing, the significance primarily lies within the inner portion of the fovea. The summarized assumptions regarding the theoretical derivation of the mapping between effective resolution and physical distance can be found in Table 1.

**Table 1.** Two regions of the receptive field on the basis of the fovea’s anatomy.

|  | Outside Fovea (OF) | Inside Fovea (IF) |
| --- | --- | --- |
| Object Size/Observation Distance | The object's image falls on both the inside and outside the fovea. This scenario mostly includes near-distance viewing. | The object's image falls on the fovea region. This scenario mostly includes far-distance (>4feet) viewing. |
| Neurophysiology | Varying visual acuity as a function of eccentricity (centered around the region of interest -ROI). This is typically modeled based on an ADF whose parameters are found through empirical means. Non-uniform cone and rod distributions. | Constant acuity inside the fovea (examples include 20/20, 20/40, etc...). Uniform cone and rod distributions. |
| Gaze Directions/frequencies | Multiple and frequent gaze direction shifts. These shifts are sufficiently separated to allow neurofusion/splining of multiple images in the higher levels of visual cortex. | Approximately a single gaze direction (or very close multiple gaze directions) and rare gaze shifts (saccades). No or only minor fusion/splining in the brain. |

In addition to these assumptions, we also assume clear atmospheric conditions that do not lead to any refraction outside the human optical system. We did not take into account the effect of rods and evaluated performance in illuminated conditions to maximize the perceived acuity. The effect of hyperacuity such as Vernier acuity is not taken into account during our experiments as they are not specifically designed for testing hyperacuity dynamics as a function of distance (For instance, there is no displacement in pixels or counters that marks the boundaries of presented faces). In fact, hyperacuity depends on the neural visual system’s power to extract small distinctions within the spatial patterns of the optical picture on the retina, whereas conventional acuity is primarily constrained by the eye’s optics and retinal mosaic. Finally, we would like to remark that hyperacuity is more related to location rather than resolution. Thus, its effect is presumed to be quite minor as far as the objectives of the study are concerned.

The central foveal region (fovea pit etc.) manifests a non-homogenous distribution of cone density, which leads to varying degrees of acuity both inside and at the periphery of the foveal region. The effective resolution values (expressed in terms of pixels) are calculated in general based on an Acuity Distribution Function (ADF) and the distances resulting from a standard geometry using angular resolution between the optotype (for example faces) and the eye’s retina (fovea pit in particular). We have estimated that an inch-long interpupillary distance ends up landing on the fovea with *∼* 2^*o*^ angular resolution (threshold angular resolution (TAR)) at around 2.1 feet distance. To make sure that the entire face (inter-temple distance of around 2 inches) will land on the fovea with high probability, we set our minimum viewing distance in our experiments between the optotype and fovea to be around 4 feet. One of the other reasons to choose our minimum distance to be a bit far from 2.1 feet is because of near-distance non-linear effects, such as refraction index of the lens and harder fusion process of different image parts of a binocular vision which may require different gaze directions to pick up necessary information.

We have originally assumed a non-uniform cone distribution inside the fovea (including the fovea pit) and the effects of rods to be minimal (as stated in Table 1). However, as we shall demonstrate that a uniform cone distribution might be a good rule of thumb under certain conditions. We have also assumed 20/20 acuity which corresponds to around 1arcmin (*∼*30 cycles per degree (cpd) - Nyquist limit max. angular resolution) inside the fovea and calculated the resulting effective resolution for distances greater than 4 feet up to 50 feet (with constant fovea size, uniform cone distribution and 1 arcmin angular resolution, corresponding to max 71.61px and min 5.73px values, respectively). We note that the model is able to accommodate different acuities as well such as 20/10 and 20/40. We finally remark that, at all distances larger than 4 feet, the faces chosen as optotypes will always land on or inside the foveal region, particularly the pit region where the photoreceptor density is the highest.

### 2.2. Modeling the Effective Resolution via Acuity Distribution: An improved Model

In order to holistically capture the peculiarities of the human acuity, it is necessary to consider peripheral acuity in addition to foveal acuity. Acuity Distribution Functions (ADF) are introduced to describe angular resolution variation over the retina as a function of gaze eccentricity (30). We have modeled the acuity degradation (or angular displacement from the center of gaze fixation) according to the following ADF function for gaze eccentricities *e > e*_0_ (31),

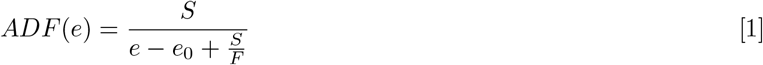

where *e*_0_ = 2^*o*^ is the average TAR assumed, *S* is the roll-off slope at the peripheral region (cpd*/*^*o*^) and typically assumed to be 65 cpd*/*^*o*^ but is subject to optimization. We would like to remind that previous empirical data shows that this parameter is typically between 55 and 80 (32). We will be adjusting this parameter based on the data we collected in our experiments. Finally, *F* is the acuity at 0 degrees gaze eccentricity (*e* = 0), which is assumed to be 30cpd for 20/20 normal vision acuity. For constant cone distribution inside the fovea, we would assume *ADF* (*e*) = *F* for *e* ≤ *e*_0_. However, constant cone distribution is recognized to be an oversimplification of the long range human visual experience (33). As a result, we recommend implementing a more concentrated cone distribution in the fovea pit for *e* ≤ *e*_0_. To achieve this, we begin by searching for the correct SF *β* that will satisfy the equation given below

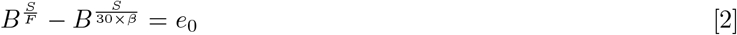

where the selection of “base *B*” depends on the available acuity. A good rule of thumb used in this study is given by *B* = 2*×* SF. Let us denote the solution to above equality as *β*^*^, then the *ADF* function for *e* ≤ *e*_0_ can be described as

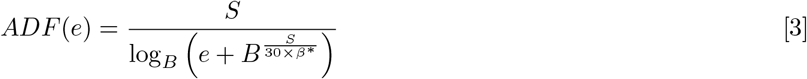

where exponentiation is introduced to model the cone packing, density of which wanes from the central pit of the fovea to the foveal avascular zone and finally to the outer periphery that extends to the macula.

Since we assume no atmospheric refraction, we hypothesize that the light emitted from an optotype can be represented by a straight line. In Fig. 2, through standard trigonometry, we observe that there is an arctangent functional relationship between the viewing distance (*x*), object/optotype size (*y*) and the viewing angle *α*. Let *d* be the inter-pupil distance and *D* be the physical distance between the optical system and the optotype. Then using the trigonometric arguments, we can estimate the gaze eccentricity in terms of degrees as

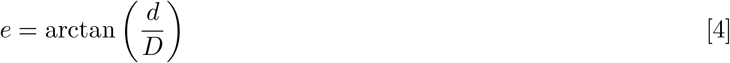

Another way to express effective resolution is in terms of dots/pixels per unit distance. According to the Nyquist-Shannon sampling theorem (34), to be able to perfectly reconstruct a full-spectrum signal, we need to sample the signal at least twice the highest frequency content of the signal unless the frequency spectrum of the signal shows sparsity. We measure frequency in terms of cpd in our context and hence the minimum sampling frequency that would allow reconstruction without *aliasing* is given by 2 *× AFR*(*e*). Accordingly, we can compute the effective resolution (*ER*) in terms of pixels as a function of distance (*D*) as follows,

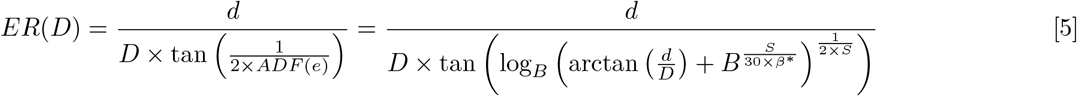

In Fig 2 (a), we have provided the acuity (cpd) as a function of eccentricity (degrees) for three different SFs, namely 20/15, 20/20 and 20/30 for both uniform and non-uniform foveal cone distributions. As can be seen the eccentricity axis is in logarithmic scale to magnify the difference of the two assumptions in the foveal region which is assumed to be below *e*_0_ = 2^*o*^ angular resolution. On the right, in Fig. 2 (b), we have plotted *ER*(*D*) function output as a function of distance in feet for 20/20 acuity and uniform foveal cone distribution for two different values of *S*. As can be seen for close viewing ranges, particularly smaller than 2.38 feet, the effective resolution is influenced by the near-range effects for practical values of *S* such as 75.

**Fig. 1.**
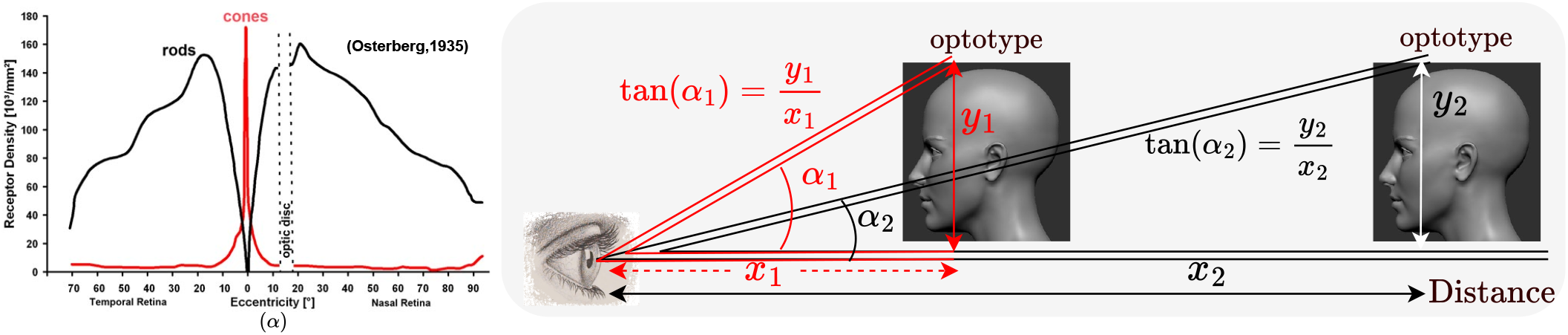
Distance and size of the optotype (face in our example) determines the angular resolution (sight angle). Through basic trigonometry, effective resolution can be computed roughly both in terms of cpd and dots/pixels per unit distance. The relationship between the viewed face image, the distance between the eye and the optotype, and the subtending angle naturally follows a triangular configuration, leading to an arctangent relationship as shown. Natural way to compute the effective resolution is to quantify the foveal area in the retina and receptors within that area responsible for processing the subtended image.

**Fig. 2.**
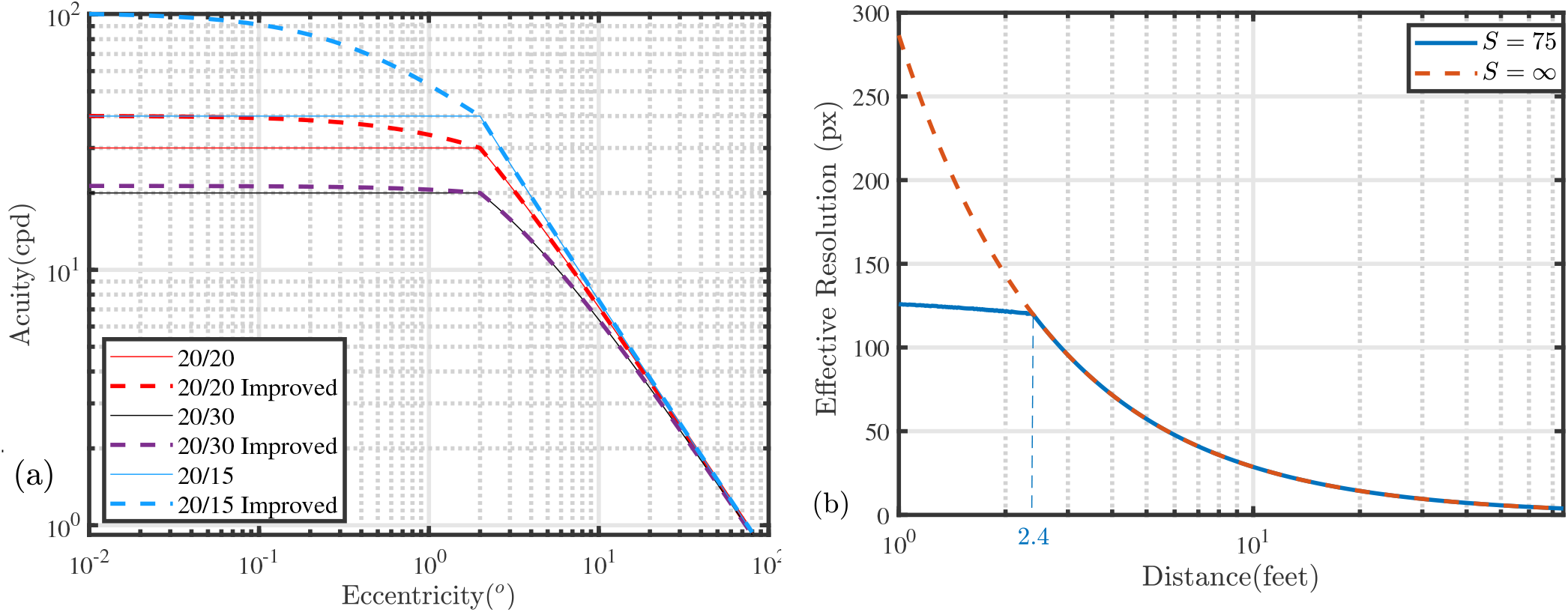
(a) Acuity (resolvable frequency) in terms of cpd as a function of eccentricity (in degrees). (b) Effective resolution function output *ER*(*D*) as a function of viewing distance *D* expressed in terms of feet and plotted in logarithmic scale. The plot also shows the effect of the parameter *S* on the effective resolution in pixels (px).

## 3. Methods

### 3.1. Subjects

Twenty normally-sighted individuals participated in this study: 11 females, 2 non-binary and 7 males (ages ranging from 18 to 59 years: An average of 28.4) from an external circle of friends/students/unfamiliar individuals who are paid to participate in the experiment. Clinical Snellen charts as well as Freiburg Visual Acuity Test (FrACT)^*^ (35) were used to evaluate each subject’s visual acuity (in logMAR units), which was shown to yield results that concur closely with traditional means. Acuity tests were measured to be around 20/20 with only minor variations between them (varying between 20/11 and 20/26). Additionally, the Pelli-Robson charts were used to evaluate each subject’s contrast sensitivity to compare with FrACT. The experiments take place within a well-lit long corridor (with a minimum range of 65 feet), wherein various data points are recorded. These include the responses indicating correctness, the time elapsed between the onset of stimulation and key pressing (response time - exposure duration), and the sequence of images presented to the subjects. Additionally, several metadata parameters, such as age, gender, experiment time, and display configuration (including luminance), are also gathered. The experiment was conducted in a controlled setting, and the participants were instructed and monitored throughout the experiment. The study conformed to the Code of Ethics of the World Medical Association (Declaration of Helsinki) and the experiments were approved by an Institutional Review Board (IRB) at the Massachusetts Institute of Technology, Cambridge, MA, USA. Each individual provided their written, informed permission prior to testing.

### 3.2. Instrumentation

We have utilized the **PsychoPy** platform for implementation, which is a free platform relying on Python-based frameworks. A 23.7-inch LG UltraFine 4K Display monitor (3840 *×* 2160 native resolution) at 60Hz refresh rate is leveraged to create 187 pixels interpupillary distance in the face images presented on the screen. The original set of images (100 identities, only male men are included) are selected from the Chicago Face Database (36). The only source of light pertaining to the presentation of stimuli in the test environment was the display monitor with adjusted luminance above 200 candela per square meter (cd/m^2^) – an average of 245 for 20 trials. The participant was positioned at one end of a well-illuminated corridor with their chin resting on a chin-rest (see also Fig. 1). A table was set up in front of them, equipped with a keyboard to enable them to react to the stimuli. The subject’s best refractive correction was used to view the stimulus display binocularly through the chin rest. The presentation display is gamma-corrected before the experiments.

### 3.3. Resampling and Low frequency representation of Images

To be able to assess the resolution of the observed object perceived by our visual system, we realize that, using standard signal processing techniques, we can resample the image plane to reduce the resolution while keeping the image size (receptive field) constant. The process of downsampling the image plane may cause high frequencies to smear and eventually lead to what is called as “aliasing” (37); hence we apply low-pass filtering (typically operationalized with a Gaussian kernel of size *σ* (sigma)) before this resampling/resizing operation. Here the 2D size of the kernel employed determines the extent of low-pass filtering and thus the *effective resolution* of the resulting image. To be able to measure human optical system performance in terms of resolution, we need to create stimuli whose resolution is reduced to a desired pixel value (accompanied with a reduction factor) using such signal processing techniques. This practice is undertaken in Appendix A to establish the relationship between *σ* and the reduction factor through optimizing the size of the Guassian kernel relative to the image size. The prominent question that arises to address the kernel size can be characterized as follows.

Given an image resampled (down-sampled and then up-sampled back to the same size using one of the interpolation methods such as *B-spline* or *bicubic*), what is the corresponding Guassian kernel size (amount of blur) that leads to the maximum similarity (such as minimum pixel-wise mean square error (MSE) or maximum structural *similarity* index (SSIM)) between the resampled version and the original image convolved with this kernel − low-pass filtered version of the image.

In our study, we have run extensive tests for three different face datasets, namely Chicago DB (36) (a total of 1203 images), Army 2m&5m (a total of 1350 images) and TinyImageNET (a total of 120,000 images) having resolutions 2444 *×* 1718, 400 *×* 330 and 64 *×* 64, respectively. We have resampled these images with varying reduction factors and investigated the means (as well as the confidence intervals) of the best Gaussian kernel size (*σ*) that minimizes the MSE or maximizes SSIM. In Fig. 3 (a), we have plotted the Gaussian blur (*σ*) parameter as a function of reduction factor whereas Fig. 3 (b) provides a few example faces from two of the databases for visualization.

**Fig. 3.**
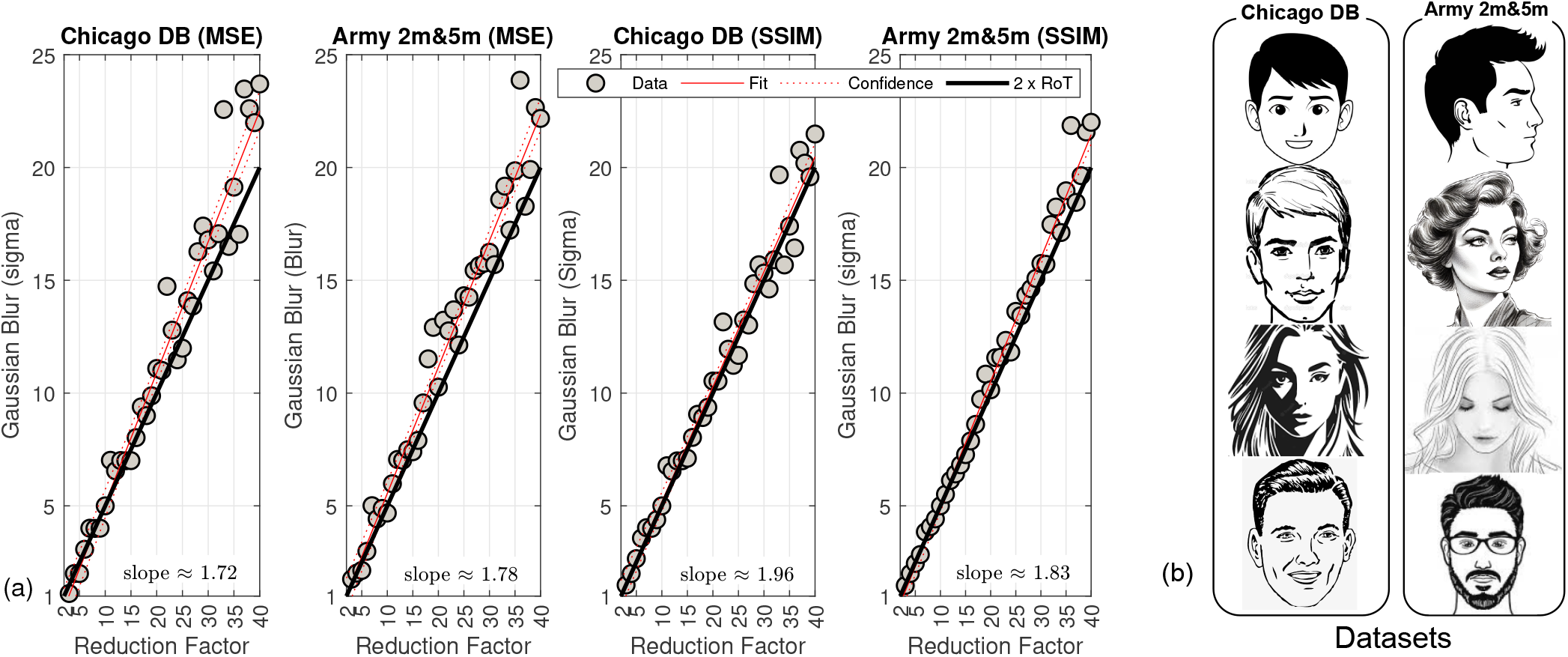
(a) This plot shows the reduction factor v.s. Gaussian Blur (kernel size expressed in terms of *σ*) for both datasets (Chicago DB and LRFID 2m&5m using similarity measures MSE and SIMM. (b) This figure shows some examples from the two datasets used to conduct this experiment. RoT: Rule of Thumb.

Particularly interesting case is Fig. 3 (a) where we clearly demonstrate that each resolution choice ended up with almost the same linear relationship between *σ* and the reduction factor. Consequently, this relationship can be captured by a simple rule of thumb (RoT); Convolving an image with a Gaussian kernel results in resolution reduction (in each dimension) by a factor of approximately 2*σ*. The black bold line in the plots shows this rule of thumb against the empirically determined red lines (and dotted line for confidence interval) for different resolutions. Furthermore, a notable observation emerges in terms of perceptual appropriateness: the SSIM, as a similarity measure, exhibits superior alignment with the anticipated 2*×* rule of thumb, in comparison to the MSE. A more theoretical exposition regarding this relationship, along with outcomes derived from the utilization of the TinyImageNET database, is provided in Appendix A.

### 3.4. Experimental Protocol Description

In our experimental design, we aimed at finding the distances at which human subjects can differentiate different quality images degraded with Gaussian blur through a task-based interaction. The monitor was placed on a cart at the furthest distance from the participant (65 feet with the physical available space. See Fig. 1). We presented 2 *×* 2 arrays of 4 images (all of the same person, only one of the four is blurred (see example shown in Fig. 4 (b)), each image size is 1360 *×* 957 pixels such that inter–pupil distance is approximately 1 inch, around 187 pixels on the display screen), and participants are given a task to identify which of the 4 images seemed blurrier than the rest (forced-choice paradigm) without time restriction. We collect responses and recorded the total time taken to generate these responses. Once completed (satisfying a certain statistical condition – Binomial test), the monitor was moved closer, to the next distance point, and the arrays of images starting from the last failed blur level were presented again. This way, we reduced the likelihood of missing any potential perceptual threshold of quality degradation. To control the bias, we used a randomized block design, in which the order of image presentation was randomized for each participant. We also leveraged a within-participant design, in which each participant saw every image, to ensure that the results were consistent across subjects. The location of the blurred image, within each array, were also randomized at every distance point. Across the arrays, there were 16 blur levels (See also Table 2) with 6 repetitions for each, with a different but fixed set of faces. The process that led to the choice of blur levels that are used in our experiment is detailed in the next subsection.

**Table 2.** One-to-one relationship between Effective Resolution, distance and Gaussian Blur (*σ*).

| Pixels | Pixels<br>(rounded) | Distance<br>(feet) | Distance<br>(inches) | Distance<br>(inches, /16)<sup>†</sup> | Reduction Factor<br>(reducing from<br>187px width) | RoT<br>Gaussian<br>Sigma<sup>‡</sup> |
| --- | --- | --- | --- | --- | --- | --- |
| 71.6197 | 72 | 4 | 48.0 | 48" 0 | 2.61 | 1.3 |
| 58.1093 | 58 | 4.93 | 59.2 | 59" 3 | 3.22 | 1.6 |
| 47.1959 | 47 | 6.07 | 72.8 | 72" 13 | 3.96 | 2.0 |
| 38.4535 | 38 | 7.45 | 89.4 | 89" 6 | 4.86 | 2.4 |
| 31.4122 | 31 | 9.12 | 109.4 | 109" 7 | 5.95 | 3.0 |
| 25.7625 | 26 | 11.12 | 133.4 | 133" 7 | 7.26 | 3.6 |
| 21.2050 | 21 | 13.51 | 162.1 | 162" 2 | 8.82 | 4.4 |
| 17.5324 | 18 | 16.34 | 196.1 | 196" 1 | 10.67 | 5.3 |
| 14.5791 | 15 | 19.65 | 235.8 | 235" 13 | 12.83 | 6.4 |
| 12.2062 | 12 | 23.47 | 281.6 | 281" 10 | 15.32 | 7.7 |
| 10.2976 | 10 | 27.82 | 333.8 | 333" 13 | 18.16 | 9.1 |
| 8.7608 | 9 | 32.7 | 392.4 | 392" 6 | 21.35 | 10.7 |
| 7.5251 | 8 | 38.07 | 456.8 | 456" 13 | 24.85 | 12.4 |
| 6.5302 | 7 | 43.87 | 526.4 | 526" 7 | 28.64 | 14.3 |
| 5.7296 | 6 | 50 | 600.0 | 600" 0 | 32.64 | 16.3 |
| 2.0000 | 2 | >50<sup>§</sup> | NA | NA | 93.50 | 46.8 |

<sup>†</sup>Distance (inches, /16)" - this is the inch distance easier to use with a ruler/measuring tape.
<sup>‡</sup>The Gaussian sigma is calculated by Factor/2 as a rule of thumb (as explained in the previous section).
<sup>§</sup>This distance was not derived from our model, but the estimation is. We added it to test our lowest boundary.

**Fig. 4.**
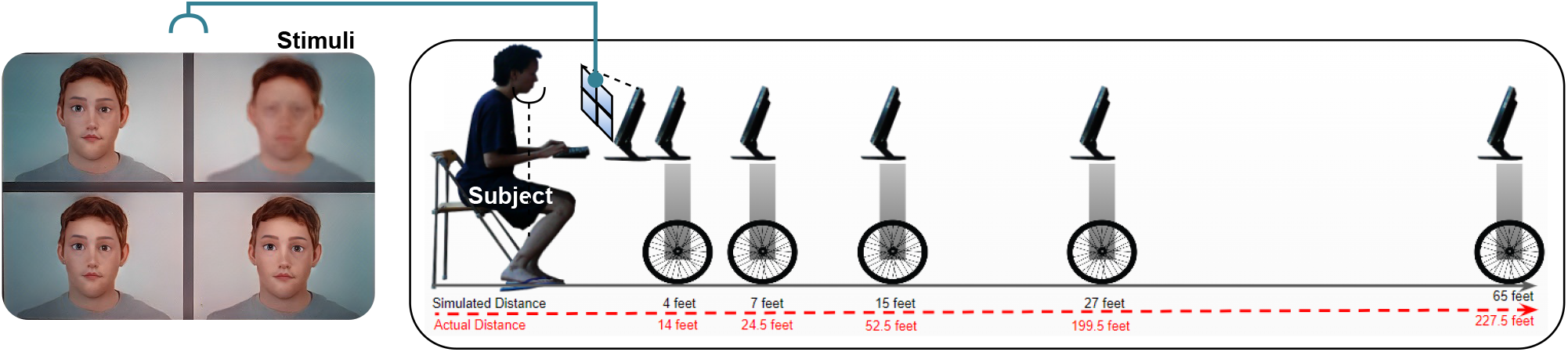
4 *×* 4 Image matrix presented to the participants where the blurred image’s location is randomized. For this example it is on the top right of the quadrant. Participant viewing the displayed images at a given distance with chin rest, completing the selection task, which can result in right or wrong decisions. The size of the face images presented on display are 3.5*×* less compared to actual average face size.

In Fig 5, we have provided the flow diagram of the experimental logic in detail. We primarily begin with the largest distance (=65 feet) and *σ* = 46.8 and decrement them based on the responses to the blocks of 6 consecutive image queries in each block. *Passes* and *Failures* are decided based on an online Binomial test which will be detailed in subsection 3.6. In case of failure, the buffer in the diagram is leveraged to store the results of the subjects’ responses from the previous block of test images which are used to decide on passing/failing on to the next distance instance based on a threshold *p* value. Once the subjects are able to pass the block, we do not go back and check again to increase our confidence. This practice is shown to dramatically reduce the experimentation time. In our experiments, the threshold *p* value is chosen to be 0.01. This implicitly implies 5 or more successful responses in the first trial would be a hard pass, whereas less than two successes would lead to a hard failure. If the subject responds to 2, 3 or 4 queries successfully in the first trial, then a second trial is executed in which a total of 8 (considering both consecutive trials which amounts to 12 trials) or more successes would eventually lead to a final pass decision, otherwise a failure will be reported.

**Fig. 5.**
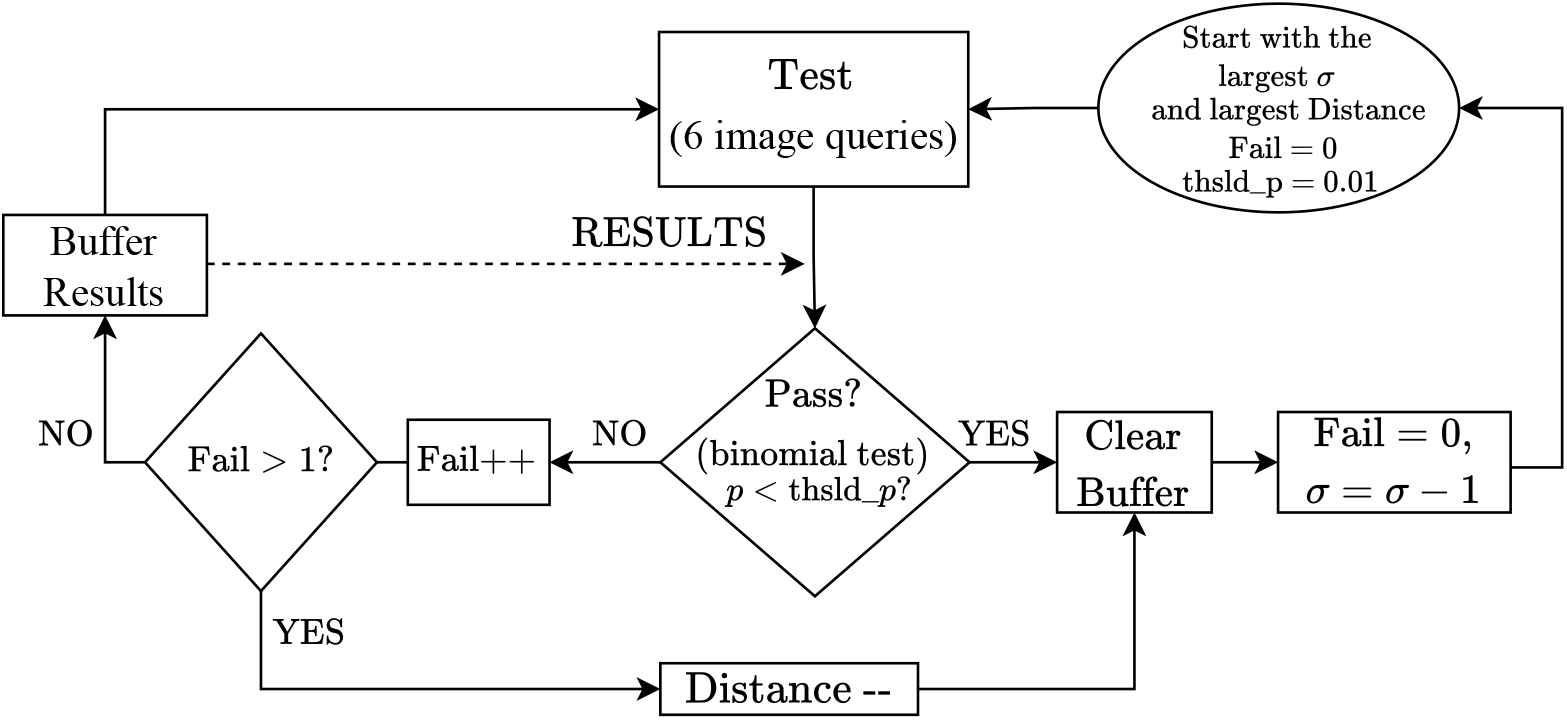
Logical flow diagram of the experiment conducted.

In Fig. 4 (a), we have shown an example 4 *×* 4 image matrix where the top-right image is degraded with blur whereas the the rest of the images are original, separated by a spacing of 0.35 inches. We also show in 4 (b) how the subject is exposed to these images with the help of a cart simulating the physical distances. For navigation between the locations of presented images, we have labeled four keys in the keyboard, namely, “e” for bottom-left, “f” for top-left, “j” for bottom-right and “o” for up-right for subjects to use and point their choice in the query image space.

### 3.5. Selection of Distance Samples

Having established the theoretical relationship captured by the ADF, we use exponential sampling (38) on the effective resolution axis to sample 15 distinct points and eventually find the corresponding 15 distance samples utilizing the inverse function *EF* ^−1^(.) to be used in our experiments. Note that we might have sampled the effective resolution axis uniformly but that has given a pretty skewed distribution of distance samples towards a shorter range (close distance) due to steep roll-off characteristics of the arctangent functional relationship. In addition, having many samples in the short range also led to the parameter of the Gaussian blur (*σ*) to be the same for some of the samples (due to close–enough sampling and rounding to one digit beyond the decimal point). To be able to have much wider coverage of distance samples for the long range, we have used exponential sampling which samples lower resolutions (farther distances) more heavily than high resolutions (shorter distances). Since in our study we are more interested in longer ranges, typically set to be higher than 4 feet, sampling long distances more heavily provides better resolution to the estimation process. The parameter of the exponential sampling, called “exponentialliness” (Φ) is set to 20 in our case (38), where the sampling function that uses this parameter can be described more specifically by

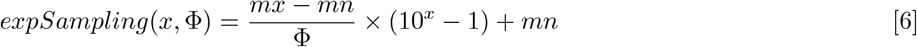

where *x* is selected from the range [0, log_10_(Φ + 1)] uniformly, *mx* is the maximum limit of the range (71.61px in our case) and *mn* is the minimum limit of the range (5.73px in our case). Note that as exponentialliness tends to unity (Φ → 1), our sampling converges to uniform sampling on the effective resolution axis with lower and upper limits [*mx, mn*]. Accordingly, we select our distances using exponential sampling and they are shown in Table 2. Note that reduction factor in resolution is assumed to be 2 as argued in the previous section.

### 3.6. Statistical Significance: Binomial Test

In a Binomial test, the null hypothesis is about finding the target response being purely a chance game, which could occur with probability 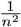 for a *n × n* array of image presentations. Accordingly, in a one-tailed binomial test, *p*-value is defined to be the probability of finding the observed number of successes (*s*) or a higher value, given that the null hypothesis is true. More specifically, we have

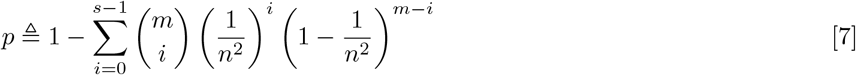

where *m* is the total number of queries in an experiment. One of the challenges to tackle for calculating the *p* value is that the average number of successes for multiple participants need not be integers. For example, in the case of multiple participants, the unbiased estimator for the accuracy of success is typically chosen to be the mean value, which might not necessarily be an integer. In order to address non-integer means, the utilization of Newton’s binomial theorem (NBT) is employed (39). When dealing with a non-integer value represented as *s*, and to ensure numerical stability, we employ the incomplete beta function, which is equivalent to NBT, for the computation of the *p* value. In other words, for any real values of *m, s, n* and 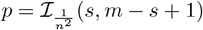, where

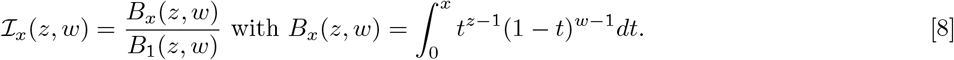

Suppose that *n* = 2, and we conduct *m* trials where we aim to achieve statistical significance at a threshold of 0.01. If we have four or more successes, the resulting *p*-value would be 0.005. However, if we only have three or more successes, the *p*-value would be 0.169. This means that we would need at least four successes to meet the required statistical significance. Similarly, if we were to perform 12 trials and aim to reach a threshold of 0.01, a minimum of 8 successes would be required. This is because if we obtain 7 successes, the resulting *p*-value would amount to 0.0143. However, with 8 successes, the *p*-value would decrease to 0.00278, surpassing the significance threshold of 0.01.

## 4. Experimental Results

### 4.1. What is the available effective resolution as a function of distance?

We begin our results by establishing the fundamental relationship between the effective resolution and physical distance (in feet) in Fig. 6 (a)-(b). Our illustration shows the mean performance of 20 subjects with 95% confidence interval. Note that based on the discrete test distance samples, we find an interval in which the effective resolution actually lies. Thus, the data points for each participant in the plot are the mean of the upper and lower resolution limits that are listed in Table 2. For instance, if the subject, at distance 133.4 inches, stumbles upon between 31.41 and 25.76 pixels of effective resolution, the data point (average) for that distance is computed to be 28.55 for that participant. The same result is complemented by two theoretical curves: One assuming a uniform–photoreceptor (Constant) fovea and the other assuming a non-linear (fovea) distribution of photoreceptors (with *S* = 55, *B* = 2.1), both based on SF 20/20. The same plot includes an alternative scale parameter on the right ordinate axis, representing the relative image size for a given distance. More specifically, the **scale** of the image that falls onto the retina is calculated based on the ratio of the effective resolution to the interpupillary distance of 187 pixels on the display screen. This way, we effectively illustrate that if one is to use scaling to replicate physical distance, the corresponding available effective resolution can be easily determined (read off) from the shared left ordinate axis of the same plot. Finally, this axis is complemented by the corresponding angular extent of the face given in units of degrees. The combined consideration of scale and angular resolution facilitates a comprehensive understanding of the impact of distant viewing on foveal vision.

**Fig. 6.**
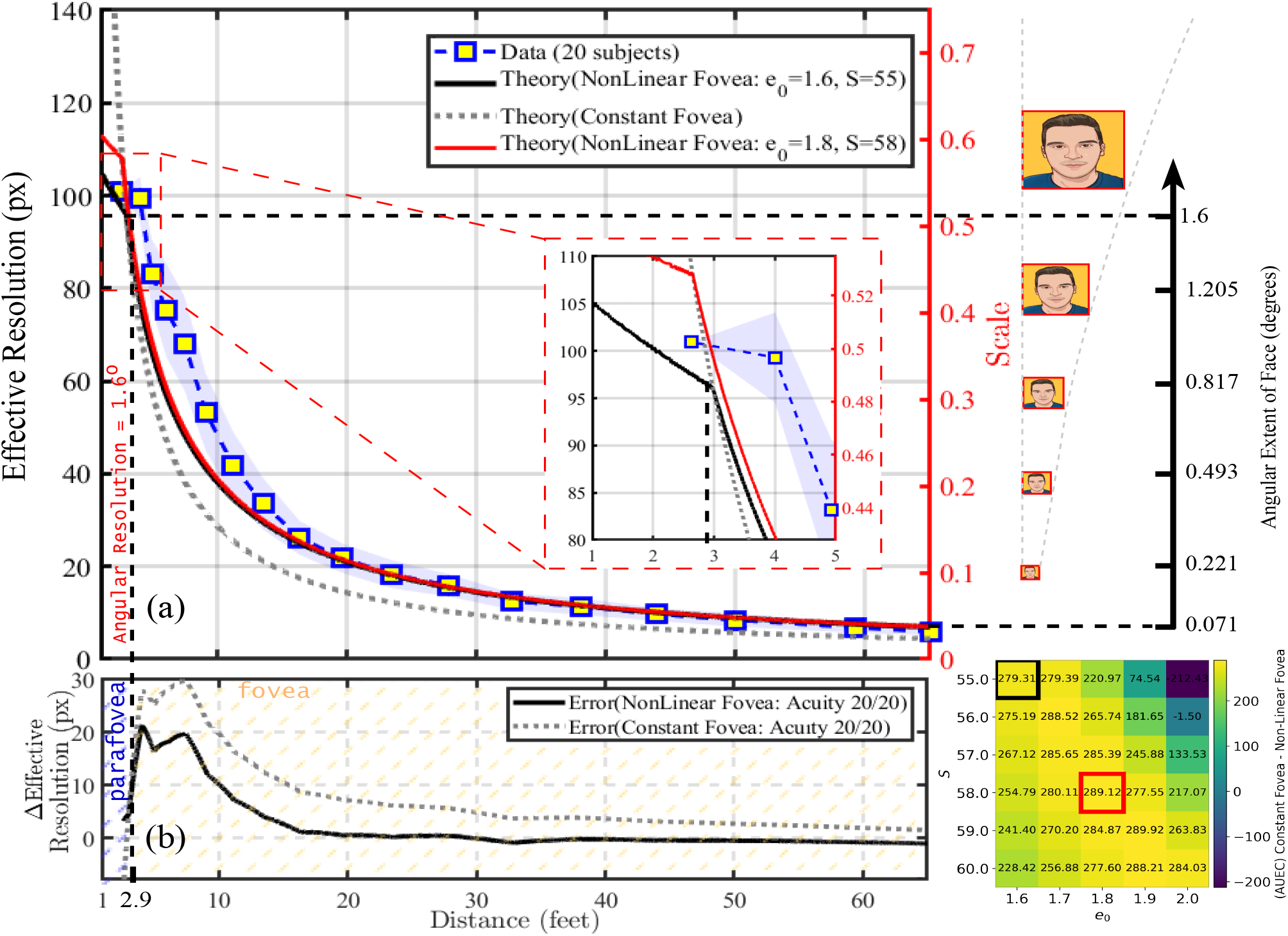
(a) The relationship between effective resolution and distance (in feet) is illustrated. The subjects’ data is represented with 95% confidence interval, accompanied by two non-linear (photoreceptor distribution) fovea models with parameters (*S* = 55, *e*_0_ = 1.6) and (*S* = 58, *e*_0_ = 1.8), both based on SF 20/20, where *e*_0_ represents the angular width of retina and *S* is the slope parameter of the ADF. The same plot includes alternative scales on the ordinate axis, one representing the corresponding image size (downscaled) that subtends the retina, the other is the angular extent (in degrees) of the face being viewed. Furthermore, we incorporated the discrapancy (the error function) observed between the gathered subject data and the two theoretical models, with a focus on both the foveal and parafoveal regions as illustrated in the subplot (b). As can be seen, the support of the error function is dominated by the foveal vision. Finally, we have included a heatmap showing the difference between the two error functions plotted in (b) for various choices of *e*_0_ and *S* in which two of these possibilities (ones that maximize the difference) are used in the main effective resolution v.s. Distance plot (subplot(a)).

One of the observations is that the proposed fovea physiology model serves as a more accurate descriptor for the functional relationship between effective resolution and distance, particularly for long ranges (see Fig. 6-(b)). However, it is noteworthy that for shorter distances, subjects surpassed the predictions of both theoretical models (maximum error), primarily due to the synergistic impact of parafoveal low spatial frequency perception and foveal acuity. This implies that further refinement of the theory may be required to provide a more comprehensive explanation for the observed outcomes.

### 4.2. Do humans adapt different strategies in resolving the blur detection task?

In order to shed light on the performance discrepancies observed among the participants in our experiments, we conducted additional analysis to investigate the following question: “Do subjects employ different strategies based on the viewing distance?”. Multiple trajectories could be explored to evaluate behavioral patterns that are most indicative of subjects’ strategies aimed at maximizing the likelihood of accuracy. Two possible trajectories we consider are (1) error patterns and (2) average latencies of true and false responses. Our hypothesis is that if subjects differ in their strategy to choose the right option, it should be reflected upon the error patterns they are generating. Thus, we focus on the former trajectory by clustering the error patterns of all participants. To accomplish this, we aggregate each subjects’ responses to create the confusion matrix as a 16-dimensional feature vector for each individual, and employ k-means clustering while optimizing the number of clusters based on the *Calinski-Harabasz* and *Silhouette* criteria^¶^. Following the clustering process, we analyze each class separately and compare their AUC performance and response latencies.

Quite remarkably, we have observed presence of two distinct groups among the participants, each exhibiting identifiable differences in their strategies. In Fig. 7 (a), we have plotted the effective resolution against simulated distance in feet for both Group 1 and Group 2. It is evident from the plot that both groups perform similarly well in the blur-detection task. To further explore the potential variations in strategies adopted by these groups, we conducted an analysis comparing the average response latency for True and False responses for each group. This relationship, depicted in Fig. 7 (b) with markings (circles and squares) representing the groups and color-coded data points representing specific distances, reveals interesting insights. Group 1, compared to Group 2, appears to allocate less time to the response generation process while achieving a similar or better AUC performance. This indicates that Group 1 employs a more efficient strategy, achieving comparable results in less time. Additionally, it is noteworthy that both groups generally spend more time on average for false responses compared to true responses which is particularly true for Group 2. This prolonged response latency for false responses may suggest the involvement of recurrent processing mechanisms, potentially engaging higher layers of cognition.

**Fig. 7.**
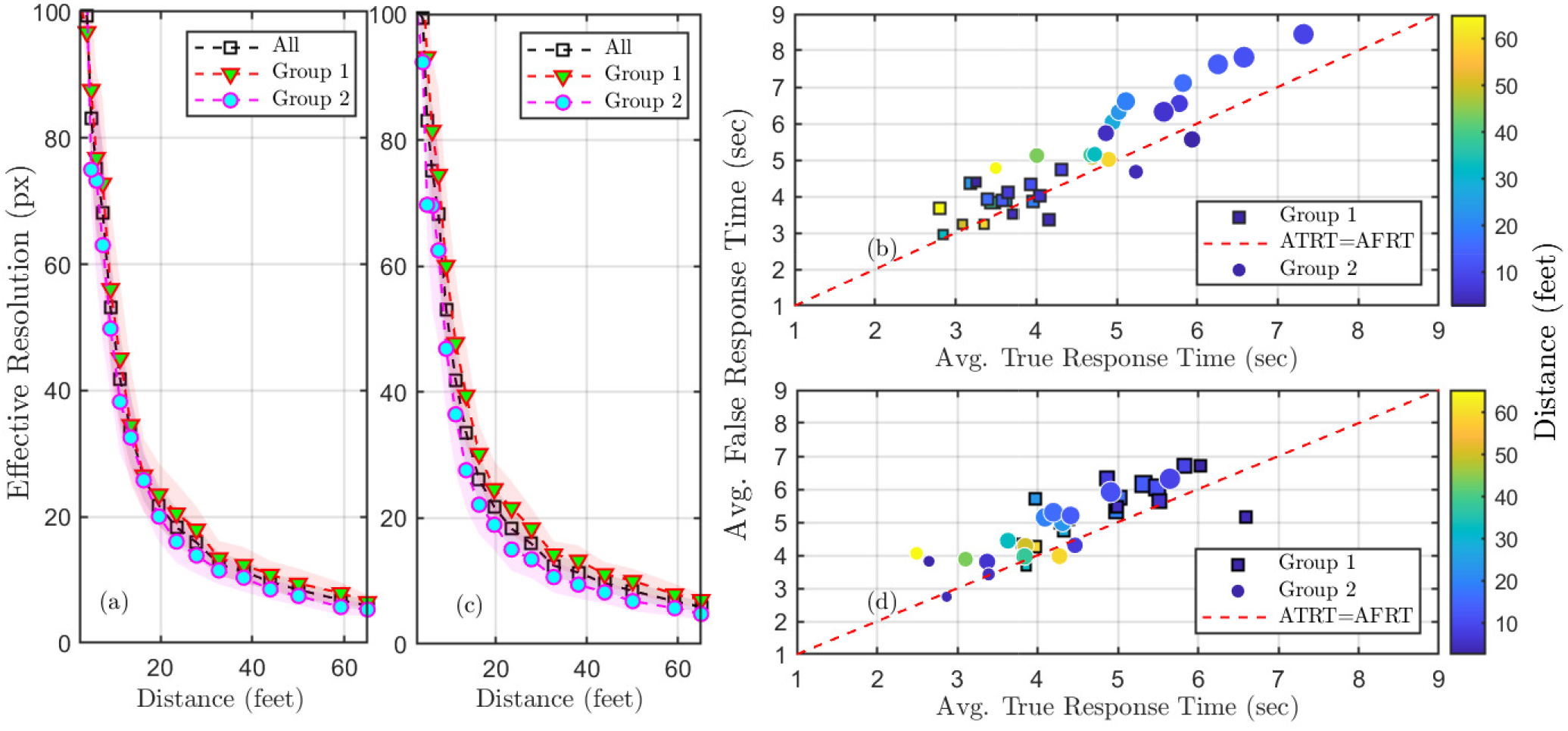
(a) The effective resolution, in relation to distance (measured in feet), for two groups of participants who were clustered based on their error patterns. (b) The average true and false response times (ATRT and AFRT, respectively), varying with distance (measured in feet), exhibit almost linear covariation, situated a little above the ATRT=AFRT line. The data is partitioned by clustering the participants’ error performances. (c) Effective resolution versus distance for two groups of participants who were clustered based on their acuity. (d) Same plot as in (b) for acuity-based clustering of participants.

### 4.3. Is acuity difference the only source of discrapancy in the performed task?

It has been observed that there is a slight distinction in the average acuity levels between the two identified groups, with mean values of -0.132 and -0.038 in units of logMAR, respectively. While it may initially appear that the performance discrepancy might arise from differences in acuity across the groups, further investigation reveals that acuity is not the sole determinant. In order to demonstrate that the discrepancy between the groups is not only due to acuity (physiological advantage), a separate clustering analysis based solely on acuity was conducted, resulting in the equal partitioning of subjects. The threshold for this partitioning turns out to be -0.085 logMAR. In Figure 7 (c) and (d), the effective resolution as a function of distance and the average response times for true and false responses are illustrated for the acuity-based groupings. It can be observed that there are more significant differences in AUC performances, which aligns with the predictive model that utilizes acuity as a parameter. Additionally, Figure 7 (d) clearly indicates that clustering based on acuity does not lead to easily distinguishable subject groups, suggesting that latency performance is not solely determined by acuity but also influenced by subsequent higher order visual as well as cognitive processes.

By employing both clustering criteria, we further conducted a comprehensive analysis, comparing AUC, Average True and False response times, contrast sensitivity, Acuity, and Age across different groups. The objective is to determine the kinds of factors besides acuity that assist these groups manifest distinctive behaviours. Our findings are presented in a star plot depicted in Fig. 8. First and foremost, it is apparent from the plot that the average time devoted to making accurate and erroneous decisions on the queries predominantly reflects the manifestation of these distinct strategies. Additionally, the star plot unveils intriguing insights, indicating that clustering based on error patterns (Fig. 8 (a)) exposes distinct strategies adopted by various age groups, regardless of their acuities and contrast sensitivities being similar. Conversely, when clustering is solely based on acuity (Fig. 8 (b)), the star plot reveals a positive correlation between higher acuity/contrast sensitivity and the improved AUC. In this clustering scenario, the groupings exhibit less distinction in terms of average group age. However, an intriguing observation emerges as the group achieving better AUC also requires more processing time, which is not the case when the clustering is performed based on error patterns. The latter can be attributed to the varying levels of confidence exhibited by the two distinct groups, which are characterized by their unique error patterns, leading to differences in the generation of true and false responses.

**Fig. 8.**
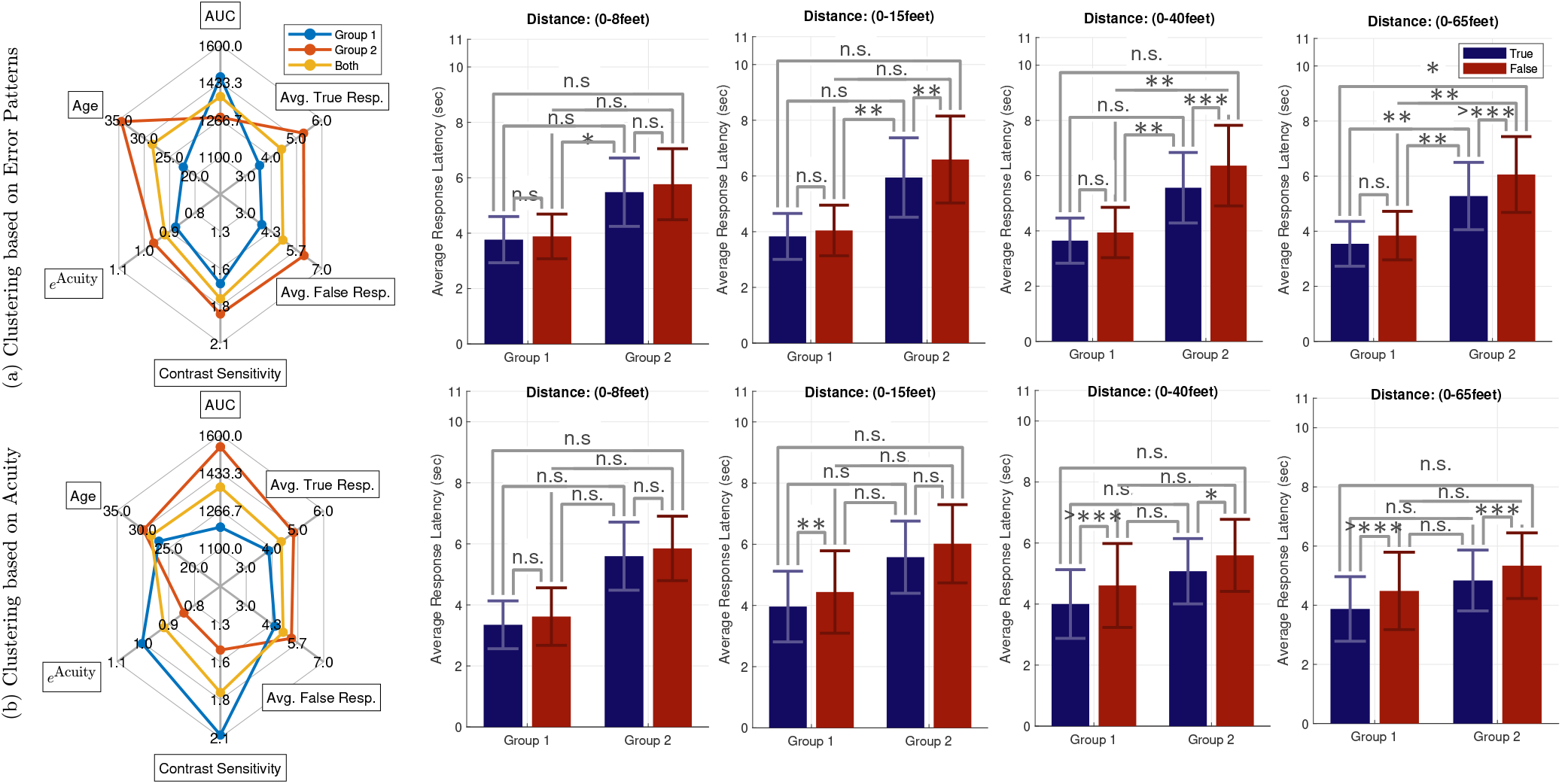
(a) The first row displays a star plot representing AUC, Avg. True Response time, Avg. False Response time, contrast sensitivity, Acuity (more accurately *e*^Acuity^: closer to 0, better the acuity), and age for the two groups identified through error pattern clustering. Additionally, the top row includes a bar plot illustrating statistical tests on average response latency for true and false responses separately using expanding windows of data (covering the first 8 feet, 15 feet, 40 feet, and 65 feet) over time. (b) The identical set of plots derived from clustering based on subjects’ acuities.

### 4.4. How do the adapted strategies change as a function of distance?

In order to examine the development of adapted strategies in relation to distance, we conducted a statistical analysis (correlation) of the average response latencies for true and false responses in both groups. Fig. 8 (a) and (b) also display bar plots indicating the average and standard error of latencies for expanding viewing ranges of 0 − 8, 0 − 15, 0 − 40, and 0 − 65 feet. First, we highlight that in our experimental setup, subjects were presented with images in a sequence where distant presentations preceded closer ones. This sequencing was deliberately chosen to reduce the potential influence of strategies tailored for short-distance viewing (better chance of identifying important cues) for long distance viewing conditions. Hence, employing an expanding window enables us to ascertain whether there exists a shift in the strategic approach to the task at hand. It is also worth noting that in our experiments, 15 feet corresponds to approximately 50 feet for real-size faces, which is considered the crucial initial recognition step marking the end of the social space (40). As observed, the differences between the groups become progressively more noticeable when clustering is performed based on error patterns, while the distinction remains consistently insignificant when clustering is based on acuity.

Another intriguing finding is that the groups with a weaker statistically significant correlation between true and false response latencies demonstrate higher performance in terms of AUC. This suggests that the failures observed may be attributed to the adoption of strategies that differ from those employed during making accurate blur detection decisions.

## 5. Discussions

As highlighted within the main content of the manuscript, our theoretical model heavily relies on several foundational assumptions grounded in the physics of light propagation, such as the assumption of rectilinear light paths, a functionally defined distribution of cones adhering to a packing density, characterized by the acuity distribution function, and the negligible influence of rods^‖^, among others. Within the framework of these assumptions, we have proposed a few improvements, notably an acuity distribution function by incorporating a non-uniform distribution of cones within the fovea, specifically within the foveal pit (foveola), and by determining the parameters of this function based on prior research (42), resulting in a more accurate alignment with the collected empirical data. However, despite these refinements, a disparity still persists between our predictive model and the experimental data, particularly evident in short-distance viewing conditions associated with the peripheral vision encompassing both non-centric foveal as well as parafoveal regions. Given that this discrepancy primarily manifests in the region nearby the periphery, we posit that it may be attributed to the non-additive effects arising from the combined influences of foveal and parafoveal vision (43). Additionally, our model does not isolate the lower/higher convergence from photoreceptor cells to ganglion cells in the foveal/parafoveal region, rather considers the combined effects of the retinal apparatus as a whole. An intriguing avenue for future research could be the design and execution of additional experiments aimed at finding the dynamics of the aforementioned combined effects by expounding on the unique transparent tissue structure (cone density drop vs. ganglion cell population change), thereby leading to the development of more refined and comprehensive mathematical frameworks.

In the previous section, we have discussed the effect of human strategies, hypothesized to reflect upon the response latency results directly. For example, we have clearly demonstrated that the differences in the errors made by the subjects cannot be explained solely by their age, acuity or contrast sensitivity. Despite the consequences of adapting different strategies are quantified, the reasons behind this visible differences are not discussed. One interesting rationale is to check as to whether there would be any difference between binocular and monocular vision as it is known that binocular summation process may improve the performance of defocus-induced blur (44). Depending on the different acuity levels of the two eyes, interocular suppression, which involves the suppression of blurred or low-contrast images in one eye by clear images in the other eye, can be observed. Later studies have shown that this supression is believed to play a role in the effectiveness of monovision correction (45), which could be helping one of the identified groups showcase a rather distinct and improved behavior.

An intriguing approach to unravel the enigmatic nature of strategies adopted by humans could involve the utilization of an eye-tracking device. By monitoring and analyzing gaze movements, it becomes possible to differentiate the saliency maps generated by the two distinct groups. This analysis would involve parameters such as the frequency and patterns of saccades, the trajectories followed during the scanning, and the location and duration of fixations, particularly within shorter viewing distances. This is due to relative size of fovea being small, requiring the eyes to continually shift their gaze to bring different portions of the image inside the foveal region. Naturally, such an endeavor would necessitate an examination of the smallest intentional saccade humans can accurately execute, as other microsaccadic movements are unlikely to play a significant role in the development of any strategy. Additionally, it would be of interest to investigate the threshold size at which humans cease their scanning movements, representing the maximum observation distance suitable for studying the human blur-detection process.

One immediate application of our mapping work is to alleviate the discrepancy between the rudimentary scaling image processing technique employed to emulate physical distance and the veritable information content that can be processed at a given distance. By translating the influence of physical distance on the observed stimuli into a quantification based on the information-bearing pixels, as opposed to a simplistic scaling approach, which is a recognized measure for neural network performance analysis, we can facilitate a more nuanced and meaningful comparison of neural network performance in relation to distance, capitalizing on having a common human visual frontend. As an illustrative demonstration, we provide an exemplar classification performance of a simple convolutional neural network on a publicly available dataset as a function of physical distance, utilizing the mapping between effective resolution/scaling and physical distance^**^. Ultimately, the objective is to enable a comprehensive comparative analysis of human recognition performance alongside computational tools, enriching our comprehension of the respective capabilities and limitations inherent in both domains. Furthermore, acquiring knowledge about human strategies is of utmost importance in constructing more effective training regimens for neural networks. This is supported by recent studies, such as (46), which align with our own findings, indicating that humans likely utilize not only shape and texture information, which may be compromised at longer viewing distances, but also higher-level “concepts” for various classification tasks (47). Therefore, conducting further investigations to uncover the underlying reasons behind these diverse strategies can assist us in training neural networks in accordance with the training protocols to which humans are developmentally exposed.

The question of whether there are developmental changes in the mapping between distance and effective resolution after acuity has stabilized is an intriguing one, particularly in view of ontogenetic development of the retina. It is notable that photosensitive ganglion cells mature very early in ontogenetic development, even before the formation of rods and the maturation of cones (41). This might suggest that there could be ongoing developmental changes in the mapping between distance and effective resolution, even after acuity has stabilized. Such changes could imply that the visual system continues to refine and adapt its mechanisms for processing visual information throughout development, perhaps in response to environmental stimuli or changing visual demands. However the specifics of this change and its eventual influence on the mapping ratios are subject to further studies. We believe that considering the evolutionary timeline of the retina may provide valuable insights into the time course of the mapping. By studying how the structure and function of the retina have evolved over time, we can gain a deeper understanding of the adaptive significance of developmental changes in visual processing and how they contribute to the functional relationship between the available resolution information and the physical distance across different species and environments.

Finally, we need to note that our study focused on testing the performance of human blur-detection task using a database of male faces. However, it remains to be explored whether the relationship between effective resolution and physical distance applies to objects and natural scenes in a general sense. While it is expected that image stimuli may not significantly impact the blur-detection task at larger viewing ranges, the situation might be little different at close ranges. Factors such as contrast differences, specialized (configural) processing, face recognition peculiarities (configuration of local and global features), and object choices can influence performance and lead to variations in decision-making processes and strategies employed. Despite conducting a computational experiment comparing the spatial frequency distribution of identities presented to different groups, no significant differences were observed. We believe that this particular outcome has the potential to serve as an initial foundation from which to embark upon a more extensive exploration on the interplay between the contextual content of stimuli and its impact on the intricacies of the blur-detection task.

## 6. Conclusions

Main objective of our study is to determine the effective resolution (a quantitative measure of available information) of face images on the retina at various viewing distances. Through a combination of theoretical considerations and experimental tasks, we observed that the relationship between viewing distance and effective resolution is more complex than initially anticipated. Our findings revealed that human performance in detecting image blur surpasses the predictions of theoretical models based on image size, cone density, and foveal extent, particularly at closer viewing ranges. This suggests the surging need for future theoretical models to account for non-uniform cone density within the fovea and better characterization of the acuity outside the fovea. By establishing a mapping between viewing distance and perceived image characteristics, our work came to have practical implications, allowing for direct comparisons of human face recognition performance under different levels of blur and viewing distances. Moreover, it facilitates systematic comparisons between human and machine vision systems by employing resolution as a common factor. Furthermore, by examining the strategies employed by humans in the blur-detection task, we aim to explore and stimulate the development of novel training strategies for neural networks, with the goal of enhancing their generalization capabilities and improving robust recognition performance. These findings highlight the importance of considering the nuanced factors affecting effective resolution and human decision making processes with varying viewing distances and pave the way for further advancements in understanding and simulating human visual perception.

## ACKNOWLEDGMENTS

This research is based upon work supported in part by the Office of the Director of National Intelligence (ODNI), Intelligence Advanced Research Projects Activity (IARPA), via [2022-21102100009]. The views and conclusions contained herein are those of the authors and should not be interpreted as necessarily representing the official policies, either expressed or implied, of ODNI, IARPA, or the U.S. Government. The U.S. Government is authorized to reproduce and distribute reprints for governmental purposes notwithstanding any copyright annotation therein.

We would like to also acknowledge Sinha Lab members for participating on the pilot versions of our experiments, research assistance Mahmoud Abdelmoneum for helping us run the experiments, and the participants of the vision sciences society (VSS) conference (2023) for sharing their comments and critics.

## Footnotes

* Available for Macintosh and Windows free of charge at http://www.michaelbach.de/fract.html

¶ Five different clusterings are conducted, and the final cluster labels are determined through Majority voting. In all instances, it was found that the ideal number of clusters is two, with each group consisting of 10 individuals, resulting in equally sized clusters.

‖ Under most lighting conditions rod and cone systems cannot operate simultaneously (41).

** This is a repository on GitHub, available for the utilization of other researchers at https://github.com/suaybarslan/effresdistCNN

